# Representation of Color, Form, and their Conjunction across the Human Ventral Visual Pathway

**DOI:** 10.1101/2020.08.28.272815

**Authors:** JohnMark Taylor, Yaoda Xu

**Author notes:** **Corresponding Author:** JohnMark Taylor, **Address:** Vision Sciences Laboratory, Department of Psychology, Harvard University, 33 Kirkland Street, Room 744, Cambridge, MA 02138, **Email:**.

## Abstract

Despite decades of neuroscience research, our understanding of the relationship between color and form processing in the primate ventral visual pathway remains incomplete. Using fMRI multivoxel pattern analysis, this study examined the coding of color with both a simple form feature (orientation) and a mid-level form feature (curvature) in human early visual areas V1 to V4, posterior and central color regions, and shape areas in ventral and lateral occipito-temporal cortex. With the exception of the central color region (which showed color but not form decoding), successful color and form decoding was found in all other regions examined, even for color and shape regions showing univariate sensitivity to one feature. That said, all regions exhibited significant feature decoding biases, with decoding from color and shape regions largely consistent with their univariate preferences. Color and form are thus represented in neither a completely distributed nor a completely modular manner, but a *biased distributed* manner. Interestingly, coding of one feature in a brain region was always tolerant to changes in the other feature, indicating relative independence of color and form coding throughout the ventral visual cortex. Although evidence for interactive coding of color and form also existed, the effect was weak and only existed for color and orientation conjunctions in early visual cortex. No evidence for interactive coding of color and curvature was found. The predominant relationship between color and form coding in the human brain appears to be one of anatomical coexistence (in a biased distributed manner), but representational independence.

## Introduction

Over the last several decades, research from psychophysics, neuropsychological case studies, neurophysiology, and neuroimaging has provided us with a wealth of knowledge regarding the representation of color and form information in the primate brain. Both color and form information have been shown to be represented and transformed across multiple levels of processing, with the relevant neural processes spanning the entire visual processing hierarchy, from the retina to higher-level ventral stream regions. Nevertheless, past studies have tended to focus on one type of feature or focus on a single brain region, leaving it unknown how the coding of color and form may change with respect to each other across the entire primate ventral processing pathway. Additionally, human fMRI studies have identified form-processing regions in lateral and ventral occipito-temporal cortex (OTC) (Malach et al., 1995; Grill-Spector et al., 1998; Kourtzi & Kanwisher, 2001; Orban et al., 2004), and both monkey neurophysiology and human fMRI studies have reported color-processing regions in ventral OTC (Hadjikhani et al., 1998; Brewer et al., 2005; Conway et al., 2007; Lafer-Sousa & Conway, 2013; Lafer-Sousa et al., 2016; Chang et al., 2017). How can we reconcile the presence of these regions showing univariate sensitivity to color or form with the fact that color and form information has been reported throughout much of the ventral visual pathway? Addressing such questions would not only further our understanding of how the primate brain represents individual features, but also shed important light on the “binding problem” of how the brain combines different visual features to construct the unified object percepts that populate our conscious visual experiences.

Characterizing the neural organization of color and form processing requires a thorough documentation of whether these two types of features are largely encoded in spatially separable modules, versus in a more distributed manner, throughout the ventral visual cortex. If processing is distributed, a further question to address is how color and form are represented together; that is, whether they are encoded in an additive/independent, versus a non-additive/interactive manner. An extreme example of non-additive coding would be a “grandmother”-type cell that only responds to a specific combination of features (e.g., one that responds to red triangles but not to other red things or non-red triangles), but such a non-additive coding scheme applies to any kind of tuning that shows an interaction effect between color and form features. Using fMRI and multi-voxel pattern analysis (MVPA), we attempt to differentiate among these possibilities and provide up-to-date documentation of the representation of color, form, and their conjunction across the human ventral visual pathway.

### Color and Form Processing Across the Visual Hierarchy

Past work has demonstrated that both color and form information is successively transformed across a series of processing stages, spanning from early visual cortex to anterior temporal lobe regions. V1 contains cells with a variety of receptive field profiles, including cells sensitive to high spatial frequencies that might subserve detailed form processing, and cells responsive to lower spatial frequencies that exhibit color tuning, which some evidence suggests may be concentrated in cytoarchitectonic regions called “blobs” (Livingstone & Hubel, 1988; Conway, 2001; Johnson et al., 2001; Johnson et al., 2009). V2 also appears to have cells with a range of different tuning profiles to color and form information, possibly with some degree of mesoscale cytoarchitectonic segregation of neurons specialized for these features (Livingstone & Hubel, 1988; Gegenfurter et al., 1996; Ts’o et al., 2001; Conway, 2010; Shapley & Hawkin, 2011). Moving onto V3, several studies have demonstrated both color and form decoding in this region (e.g., Seymour et al., 2010); however, it is more widely studied for its role in stereopsis (Adams & Zeki, 2001), and indeed some theoretical accounts of the primate color processing hierarchy omit it entirely (e.g., Conway, 2009; Conway et al., 2010).

Area V4 is an important hub for both intermediate color and form processing. Various studies suggest that it encodes mid-level form features such as curvature (Gallant et al., 1993; Gallant et al., 2000), convexity (Pasupathy & Connor, 2001), and texture (Roe et al., 2012; Pasupathy et al., 2019). V4 also encodes color information (Brewer et al., 2005; Conway et al., 2007; Brouwer & Heeger, 2009; Brouwer & Heeger, 2013; Bannert & Bartels, 2018). At least in the macaque, color-tuned cells in V4 appear to be concentrated in a handful of “glob” subregions that are each several mm in width: neurons in these “glob” regions have stronger color tuning and weaker form tuning than cells outside of them.

While color and form information clearly coexists in early visual cortex (albeit perhaps segregated by mesoscale topography), consistent with a distributed view of color and form representation, for the higher-level ventral pathway regions extending beyond V4 the distribution of neural tuning for color versus form information is more consistent with a modular view of feature representation. Regions in macaque IT cortex have famously been studied for their role in visual object processing (e.g., Tanaka, 1996; Lehky & Tanaka, 2016; DiCarlo et al., 2012), with damage to this general region leading to object processing deficits in macaques (Mishkin et al., 1983). A recent study further delineated subregions within macaque IT specialized in processing different kinds of form features (Bao et al., 2020). In human fMRI studies, high-level form processing has been linked to the lateral occipital complex (LOC) of the human brain, which comprises the lateral occipital (LO) and the ventral posterior fusiform (pFs) subregions. This region responds more to coherent than to scrambled objects, and is arguably the homolog of macaque IT cortex (Malach et al., 1995; Grill-Spector et al., 1998; Kourtzi & Kanwisher, 2001; Orban et al., 2004). Damage to this cortical region can result in loss of form perception, with spared color perception (Benson & Greenberg, 1969; Goodale & Milner, 2004). Within this broad sector, pFs but not LO appears to be invariant to mirror-image reflections (Dilks et al., 2011), and some evidence suggests that LO might be relatively more specialized for volumetric form features, with pFs being more specialized for surface and texture features (Cant & Goodale, 2007; Cavina-Pratesi et al., 2010).

Meanwhile, for color processing, a series of regions in both the macaque and human brain, extending anterior to V4, have been identified that respond more to colored than to greyscale stimuli (Hadjikhani et al., 1998; Brewer et al., 2005; Conway et al., 2007; Lafer-Sousa & Conway, 2013; Lafer-Sousa et al., 2016; Chang et al., 2017). While delineating these regions is challenging due to different naming conventions, they can be roughly grouped into posterior (overlapping with V4), central, and anterior color regions in ventral temporal cortex. The central color regions (sometimes called V8, the VO complex, or V4α; Hadjikhani et al., 1998; Bartels & Zeki, 2000; Brewer et al., 2005) have been linked to conscious color perception: they respond to color after-images (Hadjikhani et al., 1998; Humphrey et al., 1999; Morita et al., 2004), and when electrically stimulated, produce color percepts that match the color tuning of the stimulated neurons (Murphey et al., 2008; Schalk et al., 2017). The anterior color region has been shown to only respond to engaging color-involving tasks or stimuli, and not to passive viewing of colored geometric patterns (Beauchamp et al., 1999; Lafer-Sousa et al., 2016). Damage to the central and anterior color regions has been linked to neuropsychological deficits in color knowledge or color naming (reviewed in Siuda-Krzywicka & Bartolomeo, 2019) and impaired color processing with largely spared form processing (Bouvier & Engel, 2006).

The fact that there exist regions reliably showing sensitivity to color and form in univariate contrasts, and which exhibit a similar topography across participants, is consistent with a modular view of feature representation in high-level vision. Supporting this view, a human fMRI study found that color regions had no univariate preference for intact over scrambled objects (Lafer-Sousa et al., 2016). However, form information may be encoded in distributed, fine-grained activation patterns that univariate methods cannot detect (e.g., Haxby et al., 2001). It therefore remains possible that regions showing univariate sensitivity to one feature might also encode information about the other feature. Indeed, using fMRI MVPA, several human studies have found that area LO can decode color information (Bannert and Bartels, 2013; Bannert and Bartels, 2018). This is consistent with the finding that neurons in macaque IT can encode both color and form information (Komatsu & Ideura, 1993; McMahon & Olson, 2009). Likewise, neurons in macaque color regions also contained information about both color and form (Chang et al., 2017). Nevertheless, previous studies have not comprehensively documented in the human brain whether different features are represented equally strongly at the multivariate level, or whether there remains a multivariate feature coding bias consistent with a region’s univariate feature preference. Broadly, using a variety of different methods, past studies have tended to focus on particular brain regions and/or probe just one feature or the other, making it difficult to construct an overarching model of how these features are coded across the primate ventral processing pathway. A particularly important theoretical concern is reconciling the existence of regions showing univariate sensitivity for color or form with the evidence suggesting that tuning for these features might be broadly distributed throughout the ventral visual pathway. A primary goal of the present study is thus to systematically document the coding of color and form information throughout the human ventral visual processing pathway, how the relative coding strength of these two types of feature may change across brain regions, and whether there is a close correspondence in a region’s univariate and multivariate selectivity for a particular feature, using sensitive multivariate measures and well-controlled stimuli varying in their complexity. This in turn will help us understand whether a modular or a distributed view may better capture how color and form are represented together in the human brain.

### The Joint Coding of Color and Form

In addition to documenting the extent to which color and form are encoded in a modular or distributed manner, a related question to address is how these features are encoded together, whether independently, in an orthogonal manner, or in a non-additive and interactive manner.

Various lines of behavioral evidence suggest initial independent encoding of features. For example, visual search for single features is rapid, but search for feature conjunctions is slow, an observation that led Treisman and Gelade (1980) to posit *feature integration theory*: different features, like shape and color, are initially encoded on independent feature maps that can be rapidly queried, but encoding conjunctions of features requires the slow additional step of using focused attention to link the features on different maps via their shared spatial location. Consistent with this framework, “illusory conjunctions” can be induced, where the features of different objects are mismatched in conscious perception; this can occur in patients with parietal lesions, or in normal participants under conditions of divided attention (Treisman & Schmidt, 1982; Cohen & Rafal, 1991; Friedman-Hill et al., 1995). This line of research is consistent with a modular view of color and form representation, in which color and form can be independently accessed. However, it can also be consistent with a distributed view of feature representation as long as each feature can be accessed independently of the other feature. This may be achieved by having color and form represented either by commingled but distinctive neuronal populations within a given brain region or by neurons coding both features in an additive/orthogonal manner, enabling independent feature readout.

Meanwhile, several behavioral studies suggest that at least in some cases specific color and form pairings might be automatically and explicitly coded together early in processing, suggesting interactive feature coding. For example, observers can accurately report the correct pairing of color and orientation in oriented bar stimuli, even when competing color and orientation features were flickering at a high temporal frequency, suggesting that at least for certain stimuli, color and form features are automatically encoded in a conjoined format without requiring a separate, laborious attention-driven binding step (Holcombe & Cavanagh, 2001). As another example, in an illusion called the McCollough Effect, orientation-specific color aftereffects can be induced; for example, adapting to alternating red and black vertical bars will lead to a green afterimage when subsequently viewing white and black vertical bars, but not white and black horizontal bars, suggesting another case in which color and orientation might be automatically encoded in a conjunctive format (see, e.g., Stromeyer, 1969). Other studies have found further evidence of early, automatic conjunctive processing of color and form features (Victor et al., 1989; Cavanagh, 1991; Heywood et al., 1991; Barbur et al., 1994; Heywood et al., 1998; Mandelli & Kiper, 2005). Notably, these examples of early conjunctive encoding tend to involve relatively simple form features, such as orientation, leaving it unknown whether this processing format is used for the conjunction of color with more complex form features as well.

At the level of neural coding, one signature of interactive feature coding of color and form is the presence of non-additive tuning to different features in single neurons. Various studies have examined this tuning scheme. Starting with early visual cortex, a human fMRI adaptation study found that V1, V2, and V4 showed adaptation to color/orientation combinations that exceeded their predicted adaptation based on the two features individually (Engel, 2005). Another study reached a similar conclusion using multivoxel pattern analysis (MVPA; Seymour et al., 2010). Specifically, when human participants were shown pairs of alternating spiral patterns that were either red-clockwise and green-counterclockwise, or red-counterclockwise and green-clockwise, distributed activation patterns in V1 to V4 enabled decoding of these stimulus pairs from each other. This result provides evidence that there exist neural populations nonlinearly tuned to orientation and color combinations. In macaques, neurons that exhibit both color and form tuning have been found in V1 and V2 (Friedman et al. 2003), V4 (Bushnell & Pasupathy, 2012), IT (McMahon & Olson, 2009) and color regions (Chang et al., 2017). Among these macaque brain regions, non-additive coding of color and form has been reported in V4 and color regions, is largely absent in IT, and was not explicitly tested in V1 and V2. Overall, across different designs and methods, evidence for non-additive tuning has been reported in human V1 to V4, macaque V4 (and possibly macaque V1 and V2), and the macaque color regions, but not in macaque IT, or human ventral stream regions beyond V4.

A second goal of the present study is thus to examine the prevalence of nonadditive color and form coding to further our understanding of how color and form are represented together in the human brain. Thus far, this has only been studied in the context of color and orientation conjunctions in human early visual areas (Engel, 2005; Seymour et al., 2010). It remains unknown whether this coding scheme is also used for the conjunction of color with more complex form features in these brain regions. Furthermore, it remains unknown whether this coding scheme is used in higher-level ventral visual regions in the human brain.

### Present Study

In this study, we used fMRI MVPA to comprehensively chart how color is jointly represented with simple and mid-level form features across the entire known processing hierarchy for color and form in the human brain, in contrast with previous studies that primarily examined just one feature or the other, or that only examined one stage of the hierarchy. Our study will elucidate the question of whether information about color and form is confined to separate “modules” defined by univariate feature sensitivity, versus being represented in a distributed manner throughout the ventral visual processing pathway. If the latter is the case, we will further document how the *relative* strengths of color versus shape encoding might vary across brain regions, whether there is a close correspondence in a region’s univariate and multivariate selectivity for a particular feature, and whether color and form are jointly represented in an additive versus nonadditive manner. We examine these questions in the case of both simple (orientation) and mid-level (curvature) form features in two separate experiments in human early visual areas (V1 to V4), shape-selective regions, and color-selective regions (posterior and central color-sensitive regions). Our shape selective regions included lateral occipito-temporal (LOT) and ventral occipito-temporal (VOT) regions. These regions correspond to the location of LO and pFs (Malach et al., 1995; Grill-Spector et al.,1998; Kourtzi & Kanwisher, 2000), but extend further into the temporal cortex in our effort to include as many object-selective voxels as possible in occipito-temporal regions.

With the exception of the central color-sensitive region, we found that information about color and form was always co-localized in the same brain regions rather than segregated into anatomically distinctive regions in a modular fashion. This is even true for color and shape regions defined based on their univariate sensitivity to one feature. This cautions against treating regions as feature-specific “modules” based on univariate activation profiles. That said, preference of color and form information varied broadly across the regions, with preference obtained by univariate and multivariate measures by and large agreeing with each other. Color and form features are thus represented in the human brain in neither a completely distributed nor a completely modular manner, but in a biased distributed manner. Further analysis revealed that, for every region examined, coding of both color and form was tolerant to changes in the other feature. With the exception of simple form features in early visual cortex, color and form were jointly encoded in an additive and orthogonal manner throughout the human ventral visual cortex. Even for the simple form features in early visual cortex in which interactive coding of color and form did exist, the effect of interactive coding was relatively weak, suggesting that even in early visual cortex, color and simple form features are likely represented predominantly in an independent and orthogonal manner. Overall, these results provide an updated view of how color and form may be represented together in human ventral visual cortex.

## Materials and Methods

### Participants

Experiment 1 included 12 healthy, right-handed adults (7 females, between 25 and 34 years old, average age 30.6 years old) with normal color vision and normal or corrected to normal visual acuity. Experiment 2 included 13 healthy adults (7 females, between 25 and 34 years old, average age 28.7 years old). Four participants partook in both experiments. All participants gave informed consent prior to the experiments and received payment. The experiments were approved by the Committee on the Use of Human Subjects at Harvard University.

### Stimuli

#### Experiment 1: Colored spirals

Stimulus design and experimental design were adapted from Seymour et al. (2010). Participants viewed colored spiral stimuli that varied by color—red or green—and orientation—clockwise (CW) or counterclockwise (CCW)—resulting in four different kinds of spirals (Figure 1). Spirals were presented on a black background.

**Figure 1.**
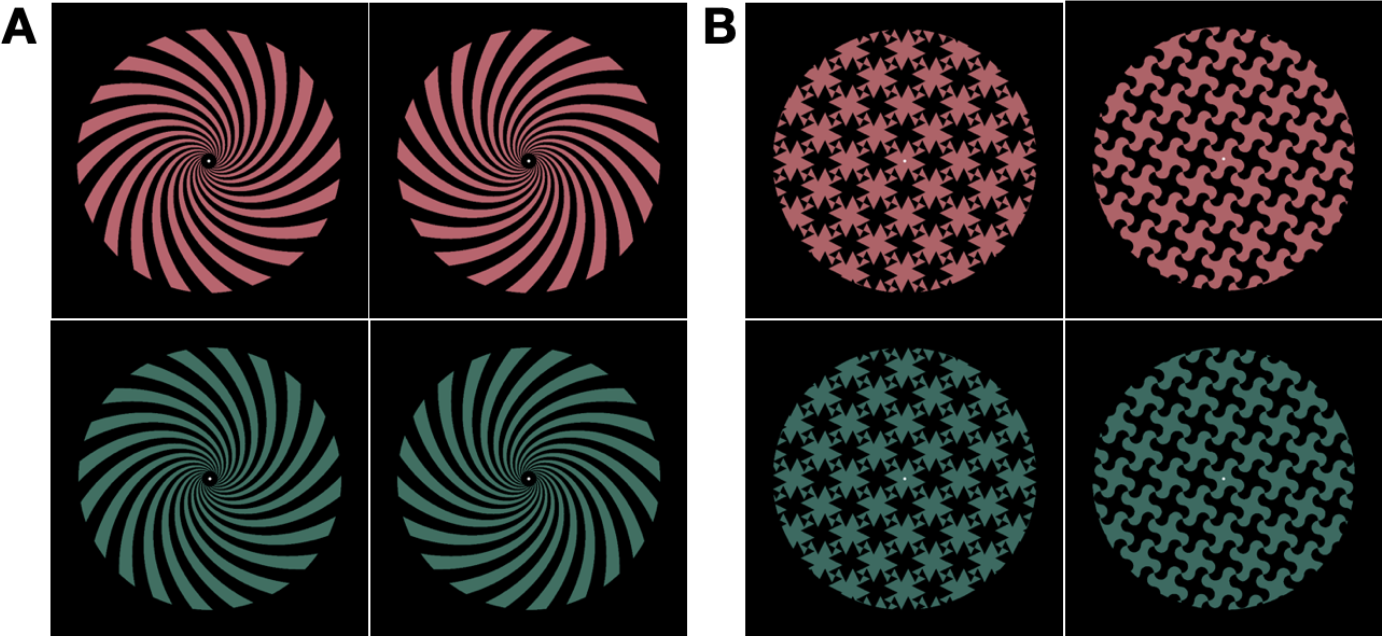
Stimuli used in the two experiments. **(A)** In Experiment 1, logarithmic spiral stimuli were shown that could be oriented clockwise or counterclockwise, and colored red or green. These spirals have the property that their arms are a fixed angle from radius at all points, ensuring that gross radial biases in cortical retinotopic maps could not drive decoder performance. These spirals were either presented in single-conjunction blocks, where a single stimulus was presented for the entire block with its phase alternating once per second, or in double-conjunction blocks, where stimuli varying with respect to both features (either RedCW/GreenCCW or RedCCW/GreenCW) alternated once per second within a block. For the double conjunction blocks, the phase of each stimulus changed every presentation, and the starting stimulus and phase were counterbalanced across blocks, such that there were four different orders for each of the two conditions. **(B)** In Experiment 2, spiky and curvy tessellation stimuli were used, with the same colors as Experiment 1. These stimuli were presented in the same manner as the spiral stimuli. For both the spiral and tessellation stimuli, the phase alternation ensures that the overall retinotopic footprint of the different form conditions are equated over the course of each block, such that this could not serve as a confound for form decoding. Additionally, for the double-conjunction blocks, the phase alternation ensures that each pixel takes on red, green, and black values a matched number of times between the two block conditions, both across the time course of a given stimulus order, and at any given timepoint for the four possible stimulus orders per block condition, ensuring that pixel-level information could not drive decoding between the two conditions.

The spirals used were *logarithmic spirals*, defined by the formula r=ae^bθ^, which have the property that the angle between the radius of the spiral and an arm of the spiral at any point is fixed, in this case at 45 degrees. This property ensures that there is a constant relationship between the location of an edge of a spiral arm in visual space and the radial component of its angle, as would not be the case if oriented gratings were used (for example, a horizontal oriented grating would have a maximal radial component along the horizontal midline, and minimal radial component along the vertical midline). This constraint accounts for the known *radial bias* in early visual cortex, in which radial orientations are preferentially represented in early visual topographic maps (e.g., zones of cortex corresponding to the top of the visual field have an over-representation of vertically oriented angles), ensuring that successful decoding of orientation could not simply be due to activation of different sub-regions of topographic maps (Sasaki et al., 2006; Mannion et al., 2009; Seymour et al., 2010). Stimuli were generated by first drawing 40 spiral lines at evenly spaced angles from the origin according to the above formula and filling in alternating regions of the spiral with the stimulus color and the background color, black, resulting in 20 spiral arms. The spiral subtended a circular region covering 9.7 degrees of visual angle, with an internal aperture in the middle, within which a white fixation dot was displayed. As mentioned earlier, the spiral arms could be oriented either clockwise or counterclockwise. Additionally, depending on which of the spiral arms were colored and which were black, each spiral could be presented in one of two phases.

The exact spiral colors used in the experiment were generated using the following procedure. To generate initially isoluminant shades of red and green, each participant performed a flicker-adjustment procedure inside the scanner, in which a flickering checkerboard with the two colors being adjusted flashed at 30 hz, and participants adjusted the colors until the flickering sensation was minimal. Specifically, the two colors had RGB values of the form red-hue = [178, 178 - X, 89] and green-hue = [0, X, 89], where participants adjusted the “X” parameter until isoluminance was achieved. This procedure guarantees that the two colors are isoluminant and sum to neutral gray, thereby equally stimulating all chromatic channels. Participants performed ten trials of this procedure, and the average “X” value was used to produce the initial colors. However, since this procedure might theoretically have some associated imprecision, each color was presented at either +/-10% of its initially calibrated luminance value on any given run of the experiment, where the number of high-luminance and low-luminance runs was balanced across the red and green colors. This manipulation ensures that any residual between-hue luminance differences will be far smaller than the within-hue luminance differences, reducing the likelihood that luminance, rather than hue, could drive MVPA classification during analysis.

#### Experiment 2: Color tessellation patterns

For this experiment, we constructed two different tessellation stimuli, consisting either of a curvy or a spiky pattern within a circular aperture (Figure 1). These stimuli were deliberately designed so as not to resemble any real-world entities, and we decided upon a curvy versus spiky contrast because curvature is a salient mid-level visual feature, in contrast with orientation, which can be considered a lower-level visual feature (Gallant et al., 1993; Srihasam et al., 2014; Yue et al., 2014). The “phase” of the tessellation stimuli could also vary, based on whether a given region of the stimulus was currently colored or black. Exactly the same procedure as Experiment 1 was used to calibrate the colors of the two stimuli, and the stimuli subtended the same visual angle (9.7°) as in Experiment 1.

### Procedure

#### Experiment 1

Participants viewed 12s blocks of the stimuli and had to detect a 30% luminance increment or decrement using a button press (index finger for increase, middle finger for decrease). On any given block, two 500ms luminance changes were presented, one in the first half and one in the second half of the block, and never in the first or last two stimuli of the block. The number and timing of the increments and decrements within the blocks was balanced across the whole experiment, and across all stimulus conditions described below. There were 9s fixation blocks between the stimulus blocks and at the end of the run, with a 12s fixation block at the beginning of the run.

The experiment included two kinds of runs. In the *single-conjunction runs*, only a single kind of spiral (RedCW, RedCCW, GreenCW, or GreenCCW) was presented for a given block, with its phase alternating once per second. This phase alternation ensures that all conditions were equated in their retinotopic footprint over the course of each block, removing this as a possible confound in form decoding. Since two starting phases were possible, each of the four spiral types could begin on either starting phase, resulting in 8 different block types for this condition. Each run contained one instance of each of the 8 types of block, totaling 180s per run. Participants completed 12 such runs, thus viewing a total of 24 blocks of each of the four spiral types over the whole session.

In the *double-conjunction runs*, there were two block conditions: a block could either alternate between RedCW and GreenCCW, or between RedCCW and GreenCW, with the phase of each spiral type alternating at each presentation. Since each block condition could begin on either one of the two spirals in one of the two phases, there were therefore four different block types for each block condition. Due to how the spirals were constructed and how the stimuli alternated phase within each block type, every pixel took on values of red, green, and black an equal number of times both over the course of any given block and at any given timepoint for the four block types within each block condition. This ensured that pixel-level information could not drive decoding during the MVPA analysis. The stimulus timing, number of blocks, and task for these runs was otherwise identical to that of the single-conjunction runs. Participants completed 12 double-conjunction runs, and thus viewed each kind of double conjunction block 48 times.

The single-conjunction runs and double-conjunction runs alternated in sets of three (e.g., three double-conjunction runs, then three single-conjunction runs), with the type of the initial run set counterbalanced across participants.

#### Experiment 2

Exactly the same task and experimental design was used in Experiment 2 as in Experiment 1, with only the stimuli varying. Due to how the tessellation stimuli were constructed and the manner in which they alternate phase within the double conjunction blocks, they shared with the spirals the property that each pixel takes on values of red, green, and black an equal number of times over the course of the block, and at corresponding timepoints for the two block conditions across the four block types within each block condition.

#### Localizer Experiments

As regions of interest in both experiments, we included retinotopically-defined regions V1, V2, V3, and V4 in early visual cortex, and functionally-defined shape and color regions in occipitotemporal visual cortex.

To localize topographic visual field maps, we followed standard retinotopic mapping techniques (Sereno et al., 1995). A 72° polar angle wedge swept either clockwise or counterclockwise (alternating each run) across the entire screen, with a sweeping period of 36.4s and 10 cycles per run. The entire display subtended 23.4 × 17.6° of visual angle. The wedge contained a colored checkerboard pattern that flashed at 4 Hz. Participants were asked to detect a dimming in the polar angle wedge. Each participant completed 4–6 runs, each lasting 364s.

We localized two shape regions in lateral occipitotemporal (LOT) and ventral occipitotemporal (VOT) cortex, following the procedure described by Kourtzi & Kanwisher (2001), and subsequently used in several of our own lab’s studies (Vaziri-Pashkam & Xu, 2017; Vaziri-Pashkam et al., 2019). LOT and VOT approximately correspond to the locations of LO and pFs (Malach et al., 1995; Grill-Spector et al.,1998; Kourtzi & Kanwisher, 2000) but extend further into the temporal cortex in order to include as many form-selective voxels as possible in occipitotemporal regions. Specifically, in a separate scanning session from the main experiment (usually the same one as the retinotopic mapping session), participants viewed black-and-white pictures of faces, places, common objects, arrays of four objects, phase-scrambled noise, and white noise in a block design paradigm, and responded with a button press whenever the stimulus underwent a slight spatial jitter, which occurred randomly twice per block. Each block contained 20 images from the same category, and each image was presented for 750ms each, followed by a 50ms blank display, totaling 16s per block, with four blocks per stimulus category. Each run also contained a 12s fixation block at the beginning, and an 8s fixation block in the middle and end. Images subtended 9.5° of visual angle. Participants performed either two or three runs, each lasting 364s.

We also localized a series of color-sensitive regions in ventral temporal cortex, using a procedure similar to Lafer-Sousa et al., (2016). Two runs of a color localizer were presented during the main scan session, one at the middle and one at the end of the session. In these runs, participants viewed 16s blocks consisting of either colorful, highly saturated natural scene images selected from the online Places scene database (Zhou, Lapedriza, Khosla, Oliva, & Torralba, 2018) or greyscale versions of these images. Participants responded when an image jittered back and forth, which occurred twice per block. Images subtended 9.5° of visual angle. Each run contained 16 blocks, 8 for each of the two stimulus types, for a total run duration of 292s including an initial 20s fixation block, and an 8s fixation block in the middle and the end of the run.

### MRI Methods

MRI data were collected using a Siemens PRISMA 3T scanner, with a 32-channel receiver array headcoil. Participants lay on their backs inside the scanner and viewed the back-projected display through an angled mirror mounted inside the headcoil. The display was projected using an LCD projector at a refresh rate of 60 Hz and a spatial resolution of 1280×1024. An Apple Macbook Pro laptop was used to create the stimuli and collect the motor responses. Stimuli were created using Matlab and Psychtoolbox (Brainard 1997).

A high-resolution T1 -weighted structural image (1.0 x 1.0 x 1.3 mm) was obtained from each participant for surface reconstruction. All Blood-oxygen-level-dependent (BOLD) data were collected via a *T*2*-weighted echo-planar imaging (EPI) pulse sequence that employed multiband RF pulses and Simultaneous Multi-Slice (SMS) acquisition. For the two main experiments, including the color localizer runs, 69 axial slices tilted 25° towards coronal from the AC-PC line (2mm isotropic) were collected covering the whole brain (TR = 1.5s, TE = 30ms, flip angle = 75°, FOV = 208m, matrix = 104×104, SMS factor = 5). For the retinotopic mapping and LOC localizer sessions, 64 axial slices tilted 25° towards coronal from the AC-PC line (2.3mm isotropic) were collected covering the whole brain (TR = 0.65s, TE = 34.8ms, flip angle = 52°, matrix = 90×90, SMS factor = 8). Different slice prescriptions were used here for the different localizers to be consistent with the parameters used in our previous studies. Because the localizer data were projected into the volume view and then onto individual participants’ flattened cortical surface, the exact slice prescriptions used had minimal impact on the final results.

### Data Analysis

FMRI data were analyzed using FreeSurfer (surfer.nmr.mgh.harvard.edu), FsFast (Dale, Fischl, & Sereno, 1999) and in-house Python scripts. The exact same analysis pipeline was used for the two experiments, except that any analyses comparing clockwise versus counterclockwise in Experiment 1 instead compared the spiky and curvy form patterns in Experiment 2, due to the differing stimuli used. Preprocessing was performed using FsFast. All functional data was motion-corrected to the first image of the run of the experiment. Slice-timing correction was applied, but smoothing was not. A generalized linear model (GLM) with a boxcar function convolved with the canonical HRF was used to model the response of each trial, with the three motion parameters and a linear and quadratic trend used as covariates in the analysis. The first eight TRs of each run (prior to the presentation of the first stimulus) were included as nuisance regressors to remove them from further analysis. A beta value reflecting the brain response was extracted for each trial block in each voxel. ROIs were defined on the cortical surface and then projected back to native functional space for further analysis.

#### ROI Definitions

##### V1 to V4

Areas V1 through V4 were localized on each participant’s cortical surface by manually tracing the borders of these visual maps activated by the vertical meridian of visual stimulation (identified by locating the phase reversals in the phase-encoded mapping), following the procedure outlined in Sereno et al. (1995).

##### LOT and VOT

Following the procedure described by Kourtzi & Kanwisher (2001), LOT and VOT were defined as the clusters of voxels in lateral and ventral occipitotemporal cortex, respectively, that respond more to photos of real-world objects than to phase-scrambled versions of the same objects (*p* <. 001 uncorrected). These regions correspond to the location of LO and pFs (Malach et al., 1995; Grill-Spector et al.,1998; Kourtzi & Kanwisher, 2000), but extend further into the temporal cortex in our effort to include as many object-selective voxels as possible in occipito-temporal regions.

##### Ventral Stream Color Regions

Following Lafer-Sousa et al. (2016), several color regions were identified in ventral temporal cortex as clusters of voxels responding more to colored images than to greyscale versions of the same images (*p* < .001, uncorrected). Since participants had varying numbers of such regions, we divided the regions in each hemisphere into anterior, central, and posterior color regions, following Lafer-Sousa et al. (2016). We were able to identify posterior and central color regions in every hemisphere of every participant in both experiments. In Experiment 1, we were able to localize the anterior color region in both hemispheres of 7/12 participants, one hemisphere of 3/12 participants, and neither hemisphere of 2/12 participants. In Experiment 2, we were able to localize the anterior color region in both hemispheres of 8/13 participants, one hemisphere of 3/13 participants, and neither hemisphere of 2/13 participants. The inconsistency in localizing this color region was likely due to its location being close to the ear canals where large MRI susceptibility effects and signal dropoff could occur. We note that our rate of localizing this color region was similar to that of Lafer-Sousa et al. (2016), who reported that this region was found in both hemispheres of 6/13 participants, one hemisphere of 4/13 participants, and neither hemisphere of 3/13 participants. These anterior regions were generally relatively small (mean 49 voxels, std 46 voxels, min 4 voxels, max 163 voxels), precluding us from conducting meaningful decoding analyses in these regions. We thus omit them from further analysis. Figure 2 shows examples from three participants. Since retinotopically-defined area V4 has also been studied for its role in color perception, we also include V4 on this figure for comparison purposes. For reference, Supplemental Figure 1 shows these regions for all participants across both experiments, since fewer studies have examined these regions compared to the early visual and shape-sensitive regions.

**Figure 2.**
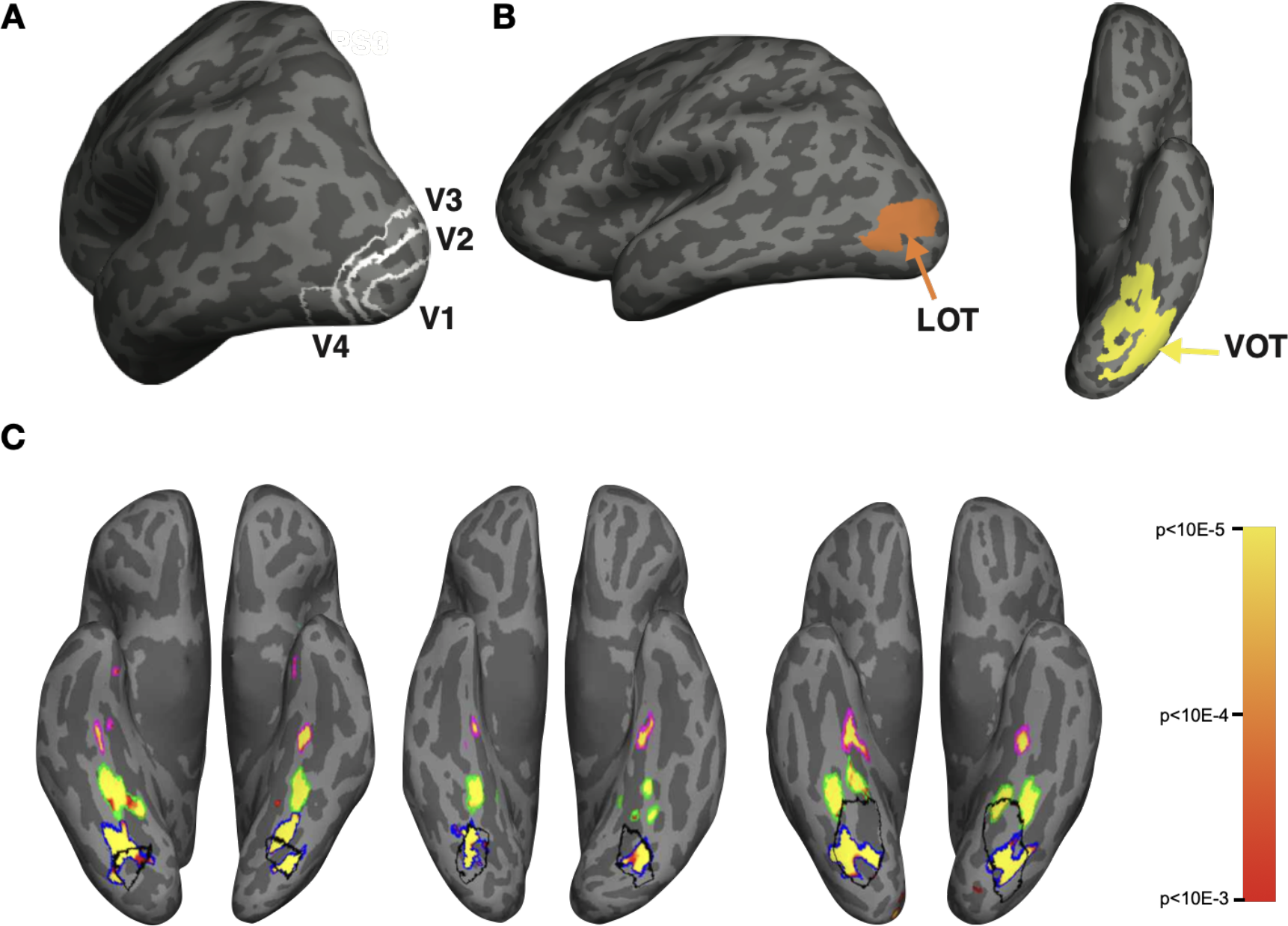
Regions of interest used in the analyses. **(A)** Lateral view of an example brain showing retinotopically-defined ROIs V1, V2, V3, and V4. **(B)** Lateral and ventral view of an example brain showing shape regions LOT and VOT. **(C)** Ventral view of an example brain showing color regions for three participants defined as higher activation for color than greyscale scenes; the posterior, central, and anterior color regions, and retinotopic V4 are shown with blue, green, magenta, and black outlines, respectively.

##### V4 and VOT *with Color Regions Removed*

We observed that the color regions overlapped with areas V4 and VOT in some cases. To document the extent to which color and form decoding in V4 and VOT might be affected by the color regions within them, we also ran several of the analyses on versions of V4 and VOT with the color-sensitive regions removed.

#### ROI Overlap Analysis

We observed that areas V4 (defined retinotopically), the posterior color region (defined using a color versus greyscale localizer), and area VOT (defined using an object versus scrambled localizer) overlapped to some degree. To quantify this overlap, we computed the pairwise percent overlap between each of these ROIs, where percent overlap was defined as the percentage of the number of overlap voxels over the averaged number of voxels for the two ROIs as we did in a previous study (Cant & Xu, 2012; see also Kung et al., 2007).

#### Multivoxel Pattern Analysis

In order to equate the number of voxels used in each ROI, the top 300 most responsive voxels in a stimulus-versus-rest GLM contrast across all the runs were selected. In addition to the ROIs described above, we also constructed an ROI for each participant consisting of the 300 most active voxels from the entire V1-V4 sector defined by the union of V1-V4. A beta value was extracted from each voxel of each ROI for every trial block. To remove response amplitude differences across stimulus conditions, trial blocks and ROIs, beta values were z-normalized across all the voxels for each trial block in each ROI. For each of the contrasts of interest (described below), these beta values were used to train and test a linear support vector machine (SVM) classifier (with regularization parameter *c* = 1), using leave-one-run-out cross-validation. T-tests were performed to compare the decoding accuracy of the various comparisons to chance (one-sample, two-tailed t-test) at the group level and, where applicable, to each other (paired t-test, two tailed), or to compare the decoding profiles across regions (two-way ANOVA). Correction for multiple comparisons was applied using the Benjamini–Hochberg procedure with false discovery rate controlled at q < 0.05 (Benjamini and Hochberg, 1995). Several different classification analyses were performed.

##### Feature Decoding

To assess the extent to which regions carried information about single features, in the single-conjunction blocks we trained and tested the classifier on color (red vs. green) and form (CW vs. CCW in Experiment 1 and curvy vs spiky in Experiment 2), where *both* values of the other feature were fed into each bin of the classifier (e.g., for color decoding, RedCW and RedCCW versus GreenCW and GreenCCW). Correction for multiple comparisons was performed within the set of comparisons done for each ROI. To test for broad trends in feature coding across the visual hierarchy, we also averaged the decoding accuracy of ROIs showing qualitatively similar response profiles via their proximity and their ordinal pattern of their feature decoding strengths over the two experiments. We additionally performed two-way ANOVAs in each ROI to examine how decoding accuracy changes based on feature (color and form), the form features used in the two experiments (orientation and curvature), and their interaction, where feature was coded as a within-subject factor and experiment coded as a between-subject factor.

Additionally, to document whether there exist any hemispheric differences in color and form coding, within each experiment we ran a matched-pairs t-test between the left and right hemisphere for both color and form coding.

Finally, to examine the extent to which feature decoding results for V4 and VOT are driven by their overlap with the color regions, we constructed ROIs consisting of V4 and VOT minus their overlap with the color regions. The same feature decoding analyses were run for these ROIs as for the other ROIs. Additionally, matched-pairs t-tests comparing the original versions of each of these ROIs to the versions with the color regions removed were run to compare both color and form decoding, pooling data across experiments to increase power.

##### Feature Cross-Decoding

To assess whether the two features were represented independently in each ROI (i.e., whether the representation of one feature was invariant across values of another feature), in the single conjunction blocks we performed cross-feature decoding in which we trained a classifier to discriminate two values of a relevant feature while the irrelevant feature was held at one value, and tested the classifier’s performance on the relevant feature when the irrelevant feature changed to the other value (e.g., train an orientation classifier on RedCW vs. RedCCW, and test orientation decoding on GreenCW vs. GreenCCW, or vice versa, with the results from the two directions averaged together). We did this for both features serving as the relevant feature. For comparison purposes, we also performed within-feature decoding, where we held the irrelevant feature constant between training and testing. This allowed us to compare the cross- and within-feature decoding using a matched number of trials. Correction for multiple comparisons was performed within the set of comparisons done for each ROI.

##### Pattern Difference MVPA

To probe for the presence of interactive color and form representation in an ROI, we ran a novel analysis to examine whether the encoding of one feature (form or color) depends on the value of the other feature. Specifically, we first took the difference between the z-normalized beta values associated with RedCW and RedCCW, and between GreenCW and GreenCCW (Figure 3). We then trained and tested an SVM (leave one run out cross-validation) on these difference vectors to examine whether the pattern differences associated with the two orientations change based on the color of the stimulus. We also performed the opposite analysis, comparing the beta value differences for the two different orientations (RedCW — GreenCW versus RedCCW — GreenCCW). The mean classification accuracies of these two directions of the analysis were then averaged. If the encoding of one feature is invariant to values of the other feature, SVM should discriminate these vectors at chance (50%); by contrast, if the encoding of one feature changes based on the other feature, the classification should be above chance. Effectively, this analysis examines whether the voxels in an ROI exhibit an interaction effect in their tuning for color and form, with the SVM classification step serving to aggregate small and potentially heterogeneous interaction effects across voxels. The same analysis was performed for the tessellation stimuli in Experiment 2, replacing CW and CCW with the spiky and curvy stimulus conditions. We performed this analysis with correction for multiple comparisons applied within each anatomical sector (early visual areas, shape regions, and color regions) for each experiment.

**Figure 3.**
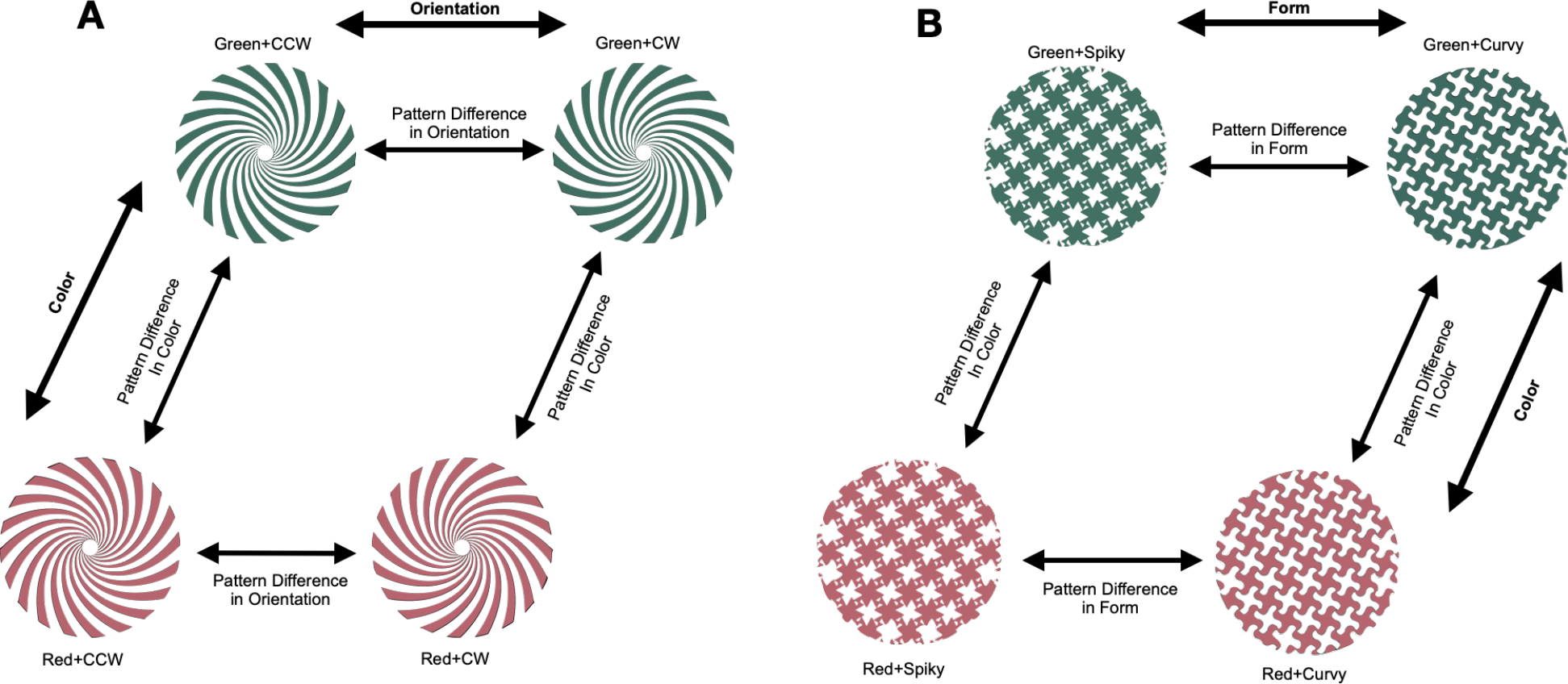
Logic of the pattern difference MVPA analysis for (A) the color and orientation spiral stimuli in Experiment 1 and (B) the color and curvature tessellation stimuli in Experiment 2. In this analysis, we examined which ROIs might code features in a manner that depends on the value of the other feature. From each ROI, we extracted and z-normalized the patterns associated with pairs of conditions matched on one feature but varying on the other, and took the difference between these patterns (e.g., GreenCCW - RedCCW). We did the same for the other value of the constant feature (e.g., GreenCW - RedCW). We then used SVM to determine whether these difference patterns were distinguishable from each other. This was done both possible ways — discriminating pattern differences in form across colors, and distinguishing pattern differences in color across the two values of each form feature — and the decoding accuracies were averaged.

We note that the information captured by this analysis is distinct from the information conveyed by feature cross-decoding. Feature cross-decoding would succeed so long as the patterns being cross-decoded end up on the correct side of the SVM decision boundary, even if the differences between the respective patterns were distinct (i.e., if main effects in feature coding far exceeded any interaction effects). By contrast, this method provides a more direct and sensitive test regarding the existence of interactive coding in the representational space.

##### Double Conjunction Decoding

As another way of examining which regions may contain interactive coding of color and form, we trained and tested the classifier on the two kinds of double conjunction blocks in each experiment (e.g., RedCW/GreenCCW and RedCCW/GreenCW). These blocks contained color and form features alternating once per second. Due to the sluggishness of the hemodynamic response, the pattern of BOLD activity present in each region would roughly constitute a superposition of the patterns associated with the two kinds of stimuli in each block. Since these two kinds of blocks both contained the two color and two form features used (e.g., red, green, clockwise, and counterclockwise), but differ in how they were conjoined, only regions encoding color and form in an interactive manner should be able to decode the two kinds of blocks from each other. The results of this analysis were compared against chance (50% decoding). As with the pattern difference MVPA analysis, we performed this analysis with correction for multiple comparisons applied within each anatomical sector (early visual areas, shape regions, and color regions) of each experiment.

## Results

Using fMRI MVPA, in the two experiments of this study, we examined the representation of simple and complex form features, color, and their conjunction in human retinotopic early visual areas (V1 to V4) and higher-level ventral regions showing univariate sensitivity to shape (LOT and VOT) and color (posterior and middle color regions). We aimed to understand whether a modular or a distributed model may better capture how color and form are represented together in the human brain, and whether these two types of features are represented in an independent/orthogonal or an interactive manner. We examined the coding of color and orientation in Experiment 1 by showing clockwise and counterclockwise spirals appearing in red and green colors, and the coding of color and curvature in Experiment 2 by showing spiky and curvy tessellations appearing in red and green colors. The phase of all stimuli alternated once per second, equating the overall stimulation across the visual field (and ruling out the possibility that any “shape” decoding could merely be due to differences in the spatial envelope of the stimuli). In some of the runs, only a single stimulus type was present in each block. FMRI pattern decoding from these runs were used to determine which brain regions contain color and/or form information and how the relative coding strength of color and form may change across the ventral visual pathway. FMRI decoding from these runs were also used in a novel measure to test the presence of interactive coding of color and form in a brain region. In the other runs, these stimuli were presented in blocks where stimuli of different forms and colors were alternated, which we analyzed using a method adapted from Seymour et al. (2010) as another metric to test the presence of interactive coding.

### ROI Overlap

Since retinotopic V4, the posterior color region, and area VOT overlap to some degree, we quantified this overlap for each pair of these ROIs. Across all the participants from both Experiments 1 and 2, V4 and the posterior color region overlapped by 40.7% +/- 2.4% (mean +/- s.e.). VOT and the posterior color region overlapped by 16.4% +/- 2.7%. VOT and V4 overlapped by 17.5% +/- 3.5%. There is thus a sizable overlap between V4 and the posterior color region, with both also overlapping slightly with VOT. Despite these overlaps, as described below, there were significant differences in how color and form were represented in these brain regions that could not be predicted by the amount of anatomical overlap. Consequently, we grouped brain regions in a later analysis by their overall functional response profile, rather than by the amount of anatomical overlap.

### Feature decoding

We used fMRI MVPA to assess the degree to which different early visual and ventral stream regions encode color and form for the two stimulus sets used in the two experiments. We both compared the decoding performance for each of the two features between the two experiments, and performed two-way ANOVAs to examine for each ROI whether decoding accuracy varies based on the feature (color or form), experiment (orientation or curvature), and their interaction. Figure 4 depicts these results.

**Figure 4.**
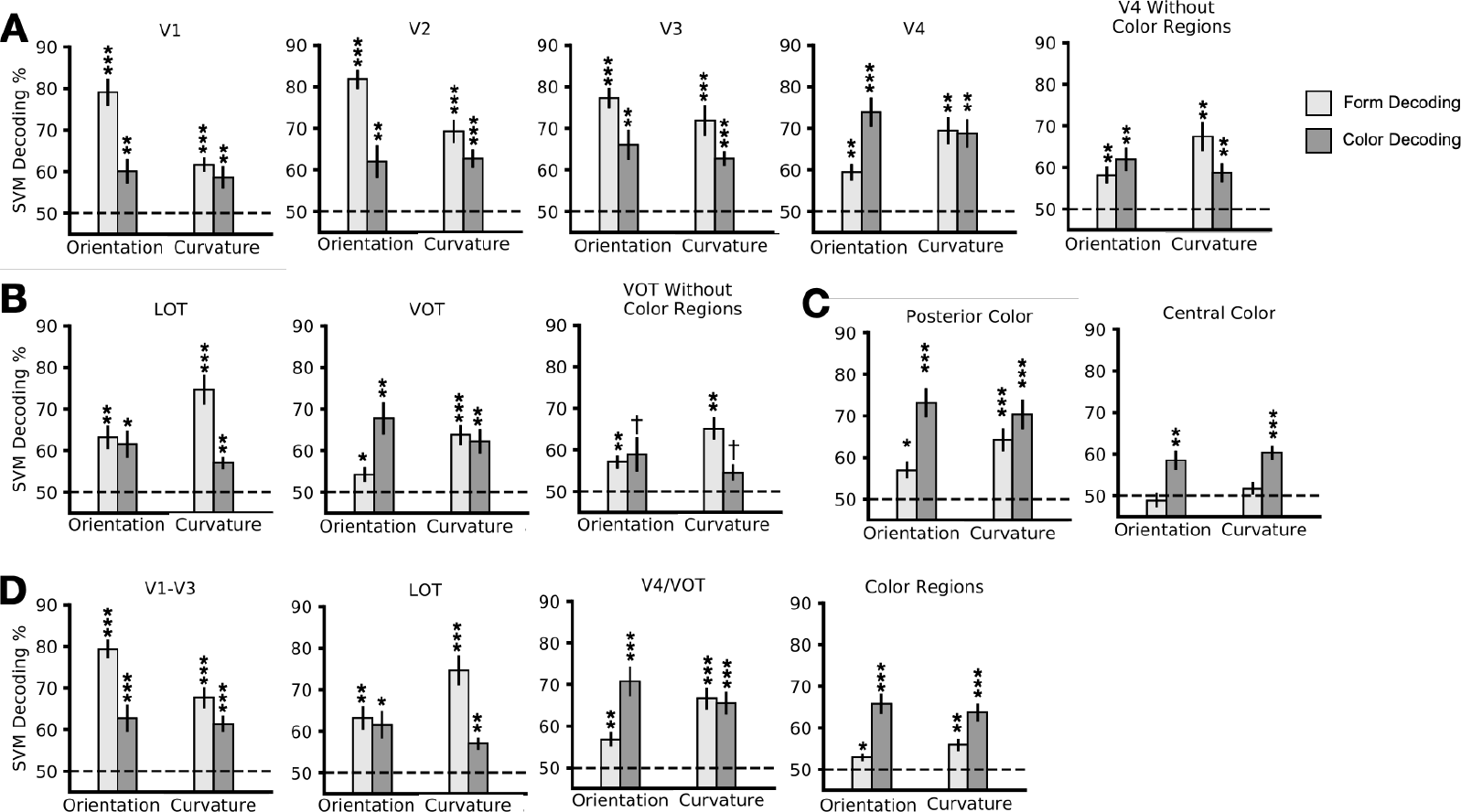
Results of color and form decoding in both experiments for **(A)** early visual areas, **(B)** shape regions, **(C)** color regions, and **(D)** sectors, which were formed based on the overall response profiles exhibited by the individual brain regions. Overall, V1 to V3 show a preference for orientation over curvature and color; VOT and V4 showed equal preference to color and curvature over orientation; the overlap of V4 and VOT with the color regions partially, but not entirely, drove color decoding in these regions; removing the color region overlap resulted in VOT showing a preference for curvature over orientation and color; LOT showed a preference for curvature over color and orientation; and lastly, the color regions showed a preference to color over either form features. * p < 0.05; ** p < 0.01; *** p < 0.001.

Beginning with the retinotopically defined early visual ROIs, pairwise t-tests against chance level performance revealed that both color and form were decodable significantly above chance in both experiments in V1 through V4 (*ts* > 3.1, *ps* < .01, two-tailed, corrected for multiple comparisons using the Benjamini–Hochberg procedure; this applies to all t-tests performed in this study, see Methods for more details). In V1 and V2, two-way ANOVAs with feature (color vs form) and experiment (Experiment 1 vs 2) as factors further showed an overall higher decoding of form than color (*Fs* > 15.9, *ps* < .001), higher decoding in Experiment 1 than 2 (*Fs* = 4.9, *ps* < .05), and an interaction between feature and experiment (*Fs* > 6.9, *ps* < .05). Pairwise comparisons within V1 and V2 showed no difference in color decoding between the two experiments (*ts* < .91, *ps* > .37) but a significantly higher form decoding in Experiment 1 than 2 (*ts* > 3.44, *ps* < .01). In V3, although an overall higher decoding for form than color was still present (*F(1,23)* = 10.8, *p* < .01), the main effect of experiment and the interaction between feature and experiment were no longer significant (*Fs* < 1.8, *ps* > .20). Pairwise comparisons revealed no difference in color or form decoding between the two experiments (*ts* < .94, *ps* >.35). In V4, there was an overall higher decoding for color than form (*F(1,23)* = 8.2, *p* < .01), with no main effect of experiment (*F(1,23)* = .461, *p* = .51), but a significant interaction between feature and experiment (*F(1,23)* = 10.5, *p* < .01). Pairwise comparisons showed higher form decoding in Experiment 2 than 1 (*t(23)* = 2.4, *p* = .04), but no difference in color decoding between the two experiments (*t(23)* = .98, *p* = .33).

Moving onto the ventral stream ROIs defined by their univariate shape sensitivity, in both LOT and VOT, decoding for both features was significantly above chance in both experiments (*ts* > 2.27, *ps* < .05). Two-way ANOVAs revealed that while there was an overall higher decoding of form than color in LOT, the opposite was true for VOT with a higher decoding of color than form (*Fs* > 7.6, *ps* < .05). In both regions, there was no main effect of experiment (*F(1,23)* < 1.1, *ps* > .31), but a significant interaction between feature and experiment (*Fs* > 9.5, *ps* <.01). Pairwise comparisons in both LOT and VOT revealed higher form decoding in Experiment 2 than 1 (*t(23)* = 2.33, *p* < .06 for LOT; and *t(23)* = 3.01, *p* = .01 for VOT), but no difference in color decoding between the two experiments (*ts > 1.07, ps* < .29).

Finally, in the ventral stream ROIs defined by their color sensitivity, for the posterior color region both features were significantly decodable in both experiments (*ts* > 2.9, *ps* < .05), with an overall higher decoding for color than form (*F(1,23)* = 14.1, *p* < .01), no main effect of experiment (*F(1,23)* = .381, *p* = .54), and a trend towards a significant interaction between feature and experiment (*F(1,23)* = 3.2, *p* = .085). Pairwise comparisons revealed no significant decoding difference for either feature between the two experiments (*ts* > .51, *ps* > .1). In the central color region, color was decodable in both experiments (*ts* > 3.5, *ps* < .01), but form was not (*ts* < 1.1, *ps* > .3), with an overall higher decoding for color than form (*F(1,23)* = 34.3, *p* < .001) but no main effect of experiment (*F(1,23)* = 1.065, *p* = .313) and no interaction between feature and experiment (*F(1,23)* = .298, *p* = .59). Pairwise comparisons revealed that feature decoding for either feature did not differ between the two experiments (*ts* < 1.2, *ps* > .48). As noted in the methods, the anterior color region identified by our localizer was very small/nonexistent in many participants and precluded us from obtaining meaningful results from this brain region in the present study.

Since V4 and VOT overlapped somewhat with the posterior color region, we performed additional analyses comparing decoding in these regions with or without the parts of these regions that overlapped with the color regions. For both ROIs (data pooled across experiments to increase power), we found no significant decrease in form decoding when the color sensitive regions were removed (*t*s < 1.81, *p*s > .08). However, in both ROIs, color decoding significantly decreased when the posterior color region was removed (*t*s > 5.15, *p*s < .001), though color decoding remained significantly above chance (*t*s > 2.89, *p*s < .01). Removing the overlapping color region from V4 and VOT also changed the relative coding strength of color and form in these regions. Notably, V4 no longer showed an overall main effect of higher color than form decoding (*F(1,23)* = .96, *p* = .33), and VOT flipped from showing higher color than form decoding to a trend for higher form than color decoding (*F(1,23)* = 3.7, *p* = .07). Other effects remained qualitatively similar as before (for both regions, no main effect of experiment, *Fs < 1.01, ps > .33*; significant interaction between feature and experiment, *Fs* > 5.15, *ps* < .03; no difference between color decoding between the two experiments, *Fs* < 1.08, *ps* > .29; and higher form decoding in Experiment 2 than 1, *t(23)* = 2.15, *p* = .08 for V4, and *t(23)* = 3.01, *p* = .01 for VOT). In other words, removing the color-sensitive voxels from VOT reversed its feature preference from stronger encoding for color to a trend for stronger encoding of form, in line with its univariate sensitivity profile.

Based on the qualitative similarity of their response profiles and their anatomical proximity, ROIs were grouped into sectors: early visual areas V1-V3, lateral visual area LOT, ventral visual areas V4/VOT, and Color Regions (including the posterior and central color regions). Decoding accuracies were averaged within sectors, and a 3-way ANOVA with feature, experiment and sector as factors verified that response profiles indeed varied significantly across these sectors (*F(3,69)* = 23.9; *p* < .001). Breaking down each of these sectors, in the V1-V3 sector color and form were significantly decodable in both experiments (*ts* > 5.5, corrected *ps* < .001), with an overall higher decoding of form than color, a main effect of experiment, and a significant interaction between feature and experiment (*Fs* > 4.6, *ps* < .05). The LOT sector only comprised area LOT, so the stats are the same as indicated above. In the V4/VOT sector, both features were significantly decodable in both experiments (*ts* > 3.40, *ps* < .01), and there was an overall higher decoding of color than form (*F(1,23)* = 9.5, *p* < .01), with no effect of experiment (*F(1,23)* = .38, *p* = .55), but a significant interaction between feature and experiment (*F(1,23)* = 13.8, *p* < .01). In the Color Regions sector, both features were significantly decodable in both experiments (*ts* > 2.5, *ps* < .05), with an overall higher decoding of color than form (*F(1,23)* = 30.1, p < .001), but no effect of experiment or interaction of feature and experiment (*Fs* < 2.79, *ps* > .11). Three-way ANOVAs (sector x feature x experiment) performed on pairs of sectors reveal significant two-way and three-way interactions involving sector, verifying that each of these sectors indeed exhibits a distinct feature encoding profile from each of the others (significant effects included: for Color Regions vs. LOT, sector x feature and 3-way interaction; for Color Regions vs. V1-V3, both sector x feature and sector x experiment, and 3-way interaction; for Color regions vs. V4/VOT, sector x feature and 3-way interaction; for LOT vs. V1-V3, sector x experiment and 3-way interaction; for LOT vs. V4/VOT, sector x feature; for V1-V3 vs. V4/VOT, both sector x feature and sector x experiment, and 3-way interaction; all *Fs* > 7.2, *ps* < .02).

Pairwise comparisons within each sector reveal no difference in color decoding between the two experiments for any sector (*Fs* < 1.54, *ps* > .227), but higher form decoding in Experiment 1 than 2 in V1-V3 (*F(1,23)* = 12.33, *p* = .002) > 5.4), lower form decoding in Experiment 1 than 2 in both V4/VOT and LOT (*Fs* > 5.4, *ps* < .03), and no difference in form decoding between the two experiments in Color Regions (*F(1,23) = 2.5, p* = .127).

We found no evidence of hemispheric differences in color coding. For form decoding, V1 and V3 showed higher decoding in the right than the left hemisphere in Experiment 1 (*ts* > 2.36, *ps* < .05); but these effects were not seen in Experiment 2 (*ts* < .60, *ps* > .56).

Overall, with the exception of the central color region, all other regions examined showed significant decoding for both color and form, even for shape and color regions showing univariate selectivity for color or form. This shows that a distributed color and form representation, rather than a strictly modular representation, is more prevalent throughout the human ventral visual cortex. Meanwhile, significant coding bias also exists in every region examined, indicating the processing of color and form is not the same in the different brain regions: even early visual areas show some feature coding preference, and in higher visual regions, such a preference appears to be largely consistent between univariate and multivariate measures.

### Feature Cross-Decoding

To understand how color and form are coded together in a brain region, we next examined the extent to which each feature is encoded in a manner that is tolerant to changes in the other feature. To do so, we performed cross-decoding and trained an SVM classifier on one feature (e.g., form) within one value of the other feature (e.g., red), and tested the classifier in the other value of the other feature (e.g., green). Additionally, to obtain a baseline measure of feature decoding with an equal amount of data for comparison purposes, we also performed within-feature decoding, and trained and tested a classifier in one feature within the same value of the other feature. Figure 5 depicts the results of these analyses. Every region that showed successful decoding of a given feature in the previous analysis also exhibited successful cross-decoding of that feature (*ts* > 2.55, *ps* < .05; color decoding in the central color regions in Experiment 1 had *ts* > 2.51, *ps* < .06). Moreover, no region showed a significant cross-decoding drop, though V1 exhibited a trend for an orientation cross-decoding drop in Experiment 1 (*t(11)* = 2.81, p = .07), and V2 exhibited a trend for a color cross-decoding drop in Experiment 2 (*t(12)* = 2.59, p = .09). These results show that regions that encode information about a feature do so in a manner that is relatively tolerant to changes in the other feature, thereby indicating some independence in representation between these two features in each region.

**Figure 5.**
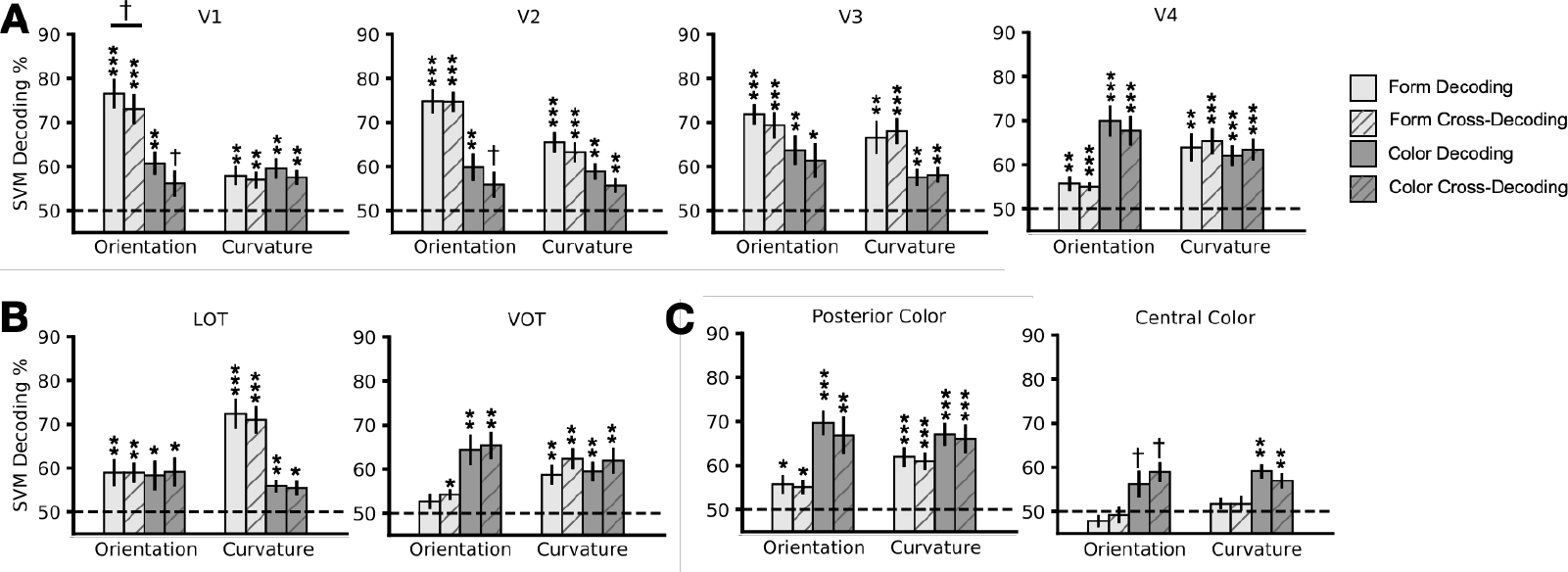
Results of feature cross-decoding analysis for **(A)** early visual areas, **(B)** shape regions, and **(C)** color regions. Solid bars show decoding accuracy for features trained and tested within the same value of the other feature (e.g., train on RedCW vs. RedCCW, test on RedCW vs. RedCCW); striped bars show decoding where training and testing for a feature is done across values of the other feature (e.g., train on RedCW vs. RedCCW, test on GreenCW vs. GreenCCW). No regions show a significant drop in cross-decoding. * p < 0.05; ** p < 0.01; *** p < 0.001; † p < 0.1.

**Figure 6.**
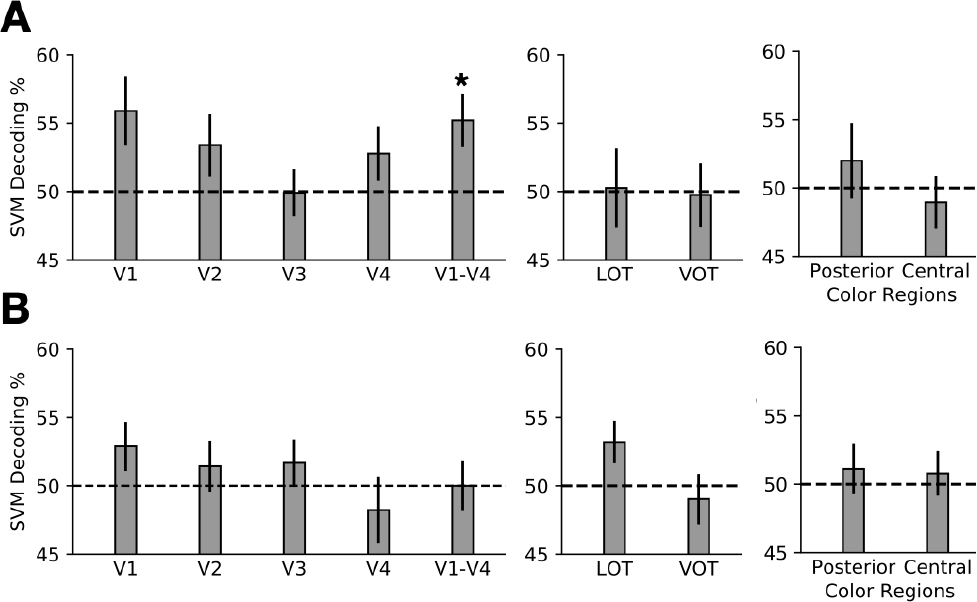
Results from pattern difference decoding analysis for **(A)** the color and orientation stimuli in Experiment 1 and **(B)** the color and curvature stimuli in Experiment 2. This analysis tested whether differences between pairs of patterns varying based on one feature (e.g., RedCW – RedCCW versus GreenCW – GreenCCW) were discriminable (see Figure 3 for a detailed explanation). Significant decoding was only obtained for the color and orientation stimuli in Experiment 1 in early visual areas. * p < 0.05.

### Pattern Synthesis Analysis

While the feature cross-decoding results show that color and form are represented in a relatively tolerant manner with respect to each other, successful cross-decoding merely requires that the test patterns lie on the same side of the SVM classification boundary as the corresponding training patterns. It does not imply a completely orthogonal representation of these two types of features (as depicted in Figure 3). To differentiate between a completely orthogonal representation and a near orthogonal representation with some level of interactive color and form conjunctive coding, we performed a *pattern difference MVPA* analysis. Specifically, we extracted two *difference vectors*, each between two stimuli that differed on the same feature dimension (e.g., one difference vector could be RedCW minus GreenCW, while the other could be RedCCW minus GreenCCW). We then tested whether these two difference vectors could be discriminated using SVM. We did this separately for both color and form and then averaged the results (see Figure 3 for a detailed illustration of this approach). If the encoding of one feature is completely independent and orthogonal to values of the other feature, then chance-level decoding is expected; by contrast, if the encoding of one feature changes based on the other feature, then above chance-level decoding is expected. This analysis essentially examines whether there is any interactive color and form coding in an ROI, with the SVM classification step serving to aggregate small interaction effects across voxels.

We found that no regions showed significant decoding (*ts* < 1.98, *ps* > .14). These results differ from those of Seymour et al. (2010) who found significant conjunction decoding with the spiral stimuli in every single early visual area examined. To increase power, besides examining each region separately, we also constructed a V1-V4 macro-ROI from the 300 most active voxels across V1-V4. With this macro-ROI, we observed significant decoding in our pattern synthesis analysis in Experiment 1 with the spiral stimuli (*t(11)* = 2.54, *p* = .03), but again failed to find significant decoding in Experiment 2 with the tessellation stimuli (*t(12)* = 1.38, *p* = .19). These results show that while interactive coding of color and form may exist in early visual areas for the simple form feature, it appears to be weak, and is absent for the complex form feature tested. Additionally, such interactive coding is largely absent in higher shape and color processing regions for both the simple and complex form features tested.

### Double Conjunction Decoding

As another way to test for the presence of interactive coding of color and form, in an independent set of data, we examined which ROIs are able to discriminate between two pairs of stimuli, where each pair has the same set of four individual features, but conjoined in different ways. Specifically, we trained a classifier to discriminate between two kinds of blocks, each consisting of alternating pairs of stimuli with different form and color features, such that the same set of four features is present in each kind of block, but combined in different ways (e.g., one kind of block alternated between RedCW spirals and GreenCCW spirals, and the other alternated between RedCCW and GreenCW spirals). If a region encodes these features in an additive, orthogonal manner, such that tuning to a feature does not depend on the value of the other feature, then patterns of activity in this region should not be able to distinguish these two kinds of block; by contrast, if they are encoded in an interactive manner, such that voxels are sensitive to particular *pairings* of color and form features, then an SVM classifier should be able to distinguish these two kinds of blocks. Figure 7 depicts the results of this conjunction decoding analysis. For the spiral stimuli in Experiment 1, across all the ROIs we examined, only area V2 showed significant interactive color and form decoding (*t(12)* = 4.1, *p* < .01). Additionally, the V1-V4 macro-ROI mentioned above also showed significant conjunction decoding (*t(12)* = 4.6, *p* < .01); we note that this same macroregion exhibited significant decoding for the pattern synthesis analysis. For the tessellation stimuli in Experiment 2, no region showed conjunction decoding (*ts* < 1.5, *ps* > .15). No additional regions became significant if no correction for multiple comparisons was made. Overall, these results largely replicate the results obtained in the pattern synthesis analysis, with interactive coding of color and form found for simple but not complex form features in early visual areas, and with such coding absent in higher visual processing regions.

**Figure 7.**
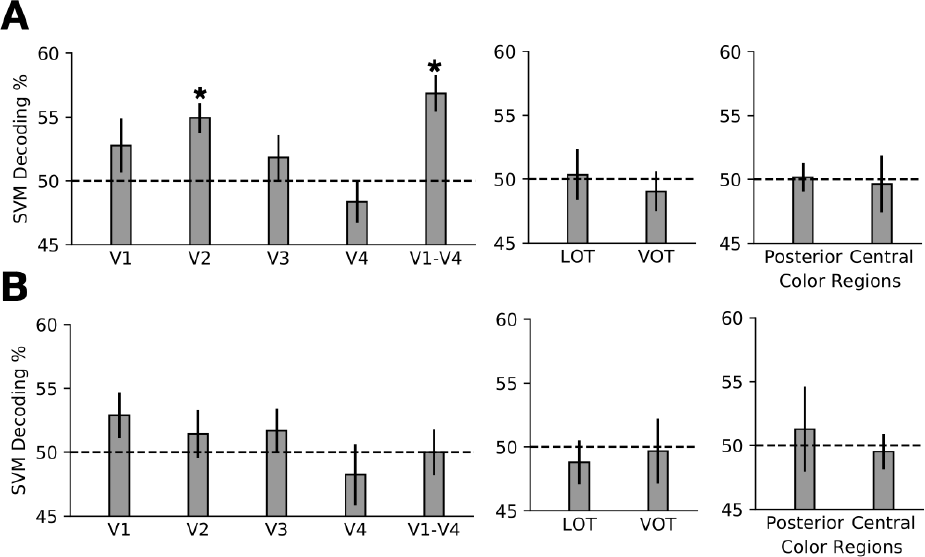
Results of double conjunction decoding analysis for **(A)** the color and orientation stimuli in Experiment 1 and **(B)** the color and curvature stimuli in Experiment 2. This analysis tested whether fMRI response patterns resulting from the display of alternating opposite-feature stimuli (e.g., RedCW/GreenCCW versus RedCCW/GreenCW) could be decoded from each other. Significant decoding was only obtained for the color and orientation stimuli in Experiment 1 in early visual areas. * p < 0.05.

These results partially replicate those from Seymour et al. (2010), who found significant conjunction decoding in every early visual ROI. It is unclear why we found weaker conjunction decoding than Seymour et al.; our experiment included more participants (12 versus 5), had more trials of the double conjunction blocks (48 versus 40), and utilized a longer ITI between blocks (9s versus 0s; Seymour et al. utilized a randomized block design with no rest periods). In order to verify that these results did not arise from a difference in our analysis methods, we re-ran the analysis with two changes to the pipeline to match the analysis of Seymour et. al. First, we included all voxels falling under *p* < .01 in a task versus rest contrast, instead of using the top 300 voxels in such a contrast. Second, instead of z-normalizing the beta values going into the analysis across voxels within each trial, we normalized the beta values of each *voxel* across all its trials. When we used the *p* < .01 activation threshold for voxel selection, we found no significant conjunction decoding in any ROI, either with within-voxel normalization or across-voxel normalization. When we selected the top 300 voxels, but used the within-voxel normalization method used by Seymour et al. (2010), we found significant conjunction coding in V2, V3, and the V1-V4 macro-ROI (*ts* > 2.6, *ps* < .05). Since the within-voxel normalization method does not equate the mean response across conditions, successful decoding in V3 for this last analysis method could have possibly arisen due to difference in mean activation across the conditions. All in all, our results partially replicate those of Seymour et al., albeit more weakly.

## Discussion

Using fMRI pattern decoding and examining color and orientation representation in Experiment 1 and color and curvature coding in Experiment 2, the present study provides a comprehensive and updated documentation of the coding of color and form information across the ventral visual processing pathway in the human brain.

### Color and form coding

Broadly, we found that color and form information is nearly always anatomically commingled in the human ventral visual pathway. This includes early visual areas V1 to V4, and higher ventral visual regions defined based on their univariate selectivity for color or shape, including the posterior color region, LOT and VOT. The only exception to this is the central color region which showed significant color decoding, but no form decoding, in both experiments. This contrasts with the results of Chang et al. (2017), who found form-tuned neurons in the macaque central color patch. We note that Chang et al. used naturalistic stimuli, rather than the artificial stimuli used in this study. It thus remains possible that the human central color patch may code both color and form features when naturalistic form features are used. We were unable to reliably localize the anterior color region in every participant here due to its location near the MRI signal dropout zone (at a rate similar to Lafer-Sousa et al., 2016). Overall, across the human ventral visual processing pathway, we found a largely distributed, rather than a modular, representation of color and form features, even in higher visual regions defined by their univariate selectivity for one feature or the other.

That said, coding preference for either feature, quantified using MVPA, varied across regions, and depended on the specific form feature tested. V1, V2, and V3 showed most sensitivity to orientation changes, and less but equal sensitivity to both curvature and color changes. These early visual areas thus showed a preference for orientation over curvature and color. VOT and V4, which greatly overlapped, showed equally strong sensitivity to color and curvature changes, but a decreased sensitivity to orientation changes. The latter could potentially be due to the mirror symmetry of the clockwise and counterclockwise spirals used, since some evidence suggests that responses in VOT may be invariant to mirror-symmetric transformations (Dilks et al., 2011). The overlap of V4 and VOT with the color regions partially, but not entirely, drove color decoding in these regions: removing the color region overlap significantly decreased color decoding in these regions, but it remained above chance. Interestingly, removing the color region overlap also resulted in VOT showing a preference for curvature over orientation and color, consistent with this region’s univariate selectivity for complex object shapes. LOT showed roughly equal sensitivity to color and orientation changes, but far greater sensitivity to curvature changes. LOT thus also showed a preference to curvature over color and orientation, consistent with its univariate selectivity for complex object shapes. Finally, the color regions showed greater sensitivity to color than either orientation or curvature changes, consistent with their univariate sensitivity to color. Thus, despite an overall distributed representation of the color and form features, even early visual areas show a feature preference, and in higher visual regions, their feature preferences are largely consistent between univariate and multivariate measures. Thus color and form features are represented in the human brain in neither a completely distributed nor a completely modular manner, but in a biased distributed manner.

We note that the color and form information encoded in different regions may play different roles in visual information processing. For example, only feature information in some regions may be directly available to conscious perception, whereas the same information in other regions could be put to other uses. This can be seen in achromatopsia patients who can perceive isoluminant, color-defined shapes (e.g., a red square on a green background), even if they cannot report the colors that define the shape (Victor et al., 1989; Heywood et al., 1991; Barbur et al., 1994; Heywood et al., 1998). Perhaps this could be one functional role of color information in form-processing regions such as LOT. Differences in how the encoded feature information is utilized by the visual system may explain the human lesion results and visual search results that on the surface support a modular view of feature processing in the human brain, even though the underlying neural representation may follow a biased distributed organization as shown in this study.

### Joint Coding of Color and Form

To understand how color and form may be represented together, we performed several additional analyses. Using a cross-decoding approach, we found that encoding of both color and form was tolerant to changes in the other feature for every ROI we examined. This indicates some independence in representation between these two features in each region. At the circuit level, such independence could be achieved by either intermingled but specialized neurons tuned to each feature, or by neurons tuned to both features but responding in a linearly additive manner. The existence of both types of neurons in various cortical regions have been reported by neurophysiological studies (as reviewed in the introduction).

However, cross-decoding success only requires that the test patterns lie on the same side of the SVM classification boundary as the corresponding training patterns. It does not preclude the co-existence of neurons with nonlinear interactive coding of color and form that exhibit tuning to particular feature conjunctions. Using a novel pattern difference MVPA analysis that can aggregate small interaction effects across voxels, we found evidence for interactive coding of color and orientation in a macro-ROI consisting of V1-V4. In a separate data set, using the double conjunction methodology developed from Seymour et al. (2010), we also found evidence for interactive coding of color and orientation in V2 and the macro-ROI consisting of areas V1-V4. Notably, we did not find interactive coding of color and orientation in higher visual areas nor did we find such coding for color and curvature in any region examined.

Although our method depends upon neural tuning being heterogeneously clustered across voxels in a way that can be detected by the spatial resolution of fMRI, the presence of strong color and form decoding across brain regions indicates strong fMRI-detectable heterogeneity in feature representation across the visual cortex. If neuronal populations exhibiting interactive coding exist, it is surprising that this heterogeneity in distribution should be absent for the interactive coding of color and simple form in higher visual regions when it is present in early visual areas, and absent for the interactive coding of color and mid-level form in all regions we examined. At the very least, if these neuronal populations do exist, we can conclude that they are distributed very differently from the other neuronal populations involved in color and form coding in the ventral visual cortex. It is also possible such neuronal populations simply do not exist, thereby avoiding the potential combinatorial explosion involved in having dedicated neurons for encoding the combination of every form and every color. It should be noted that even in early visual cortex we obtained much stronger decoding results for single features than for feature conjunctions and that cross-decoding revealed color and form coding that was tolerant to changes in the other feature. This suggests that, despite the presence of interactive color and orientation coding in early visual areas, features are still likely represented predominantly in an independent and orthogonal manner in these regions.

In their study examining color and form coding in the macaque color patches, Chang et al. (2017) found neurons that exhibited a color and form interaction effect. We note that a variety of form categories were used in that study, including faces, animals, fruits, man-made objects and geometric forms. Thus an interaction of color and form coding could reflect some category-specific effects that may not be applicable when novel form features are used, such as those used in the present study. Further studies are needed to test whether such an interaction of color and form coding still exists in color regions when novel geometric forms are used.

Treisman and colleagues have famously argued that independently coded features can be conjoined via their shared location (Treisman & Gelade, 1980). One proposed neural mechanism for achieving this has been long-range synchronized firings between neurons corresponding to different features of the same object at the same spatial location (Singer, 1999), with the posterior parietal cortex (PPC) serving a critical role in mediating this process (Robertson, 2003) as damage to PPC can result in feature binding deficits (Cohen & Rafal, 1991; Friedman-Hill et al., 1995). However it is unclear how such a code would be generated and read out, and the wiring patterns and temporal firing precision of neurons between brain regions may be insufficient to implement this code (Shadlen & Movshon, 1999). Nevertheless, binding through a shared location via a neural mechanism other than synchrony is still possible. With the exception of the color regions beyond V4, detailed spatial representation has been demonstrated for all the other ROIs tested (though there remains a possibility that the central color regions overlap with the retinotopic maps VO1 or VO2; see Brewer et al., 2005; Larsson & Heeger, 2006; Wandell & Winawer, 2011). The co-existence of color and form representation within most ROIs we examined, together with the co-existence of a detailed spatial map, could facilitate a binding by location mechanism at the local level without evoking long-range couplings between brain regions through neural synchrony, thereby serving as a potential binding mechanism (see also Di Lollo, 2012).

If binding may be achieved at the local level, then why might damage to PPC result in feature binding deficits (Cohen & Rafal, 1991; Friedman-Hill et al., 1995)? Recent work has shown the important role of PPC in visual working memory and attention-related visual processing (e.g., Bettencourt & Xu, 2016; Vaziri-Pashkam & Xu, 2017; see also Xu, 2017 & 2018a). It has been proposed that PPC may constitute a second visual system in the primate brain and participate in the adaptive aspect of visual processing by extracting and retaining task relevant visual information (Xu, 2018b). Thus a failure to perceive a correct feature conjunction due to PPC damage could be due to a failure to extract and tag the correct feature information from ventral regions for task-related visual processing. Although more research is needed to fully explore this possibility, the present study charts the anatomical layout and coding scheme of the ventral stream feature representations over which any putative parietal mechanism involved in feature binding must operate.

In this study, we compared the magnitude of color and form decoding. It could be argued that this comparison depended on the amount of variation we introduced within each feature. For example, by reducing or increasing the difference between the two colors we examined, we could change the magnitude of color decoding and shift the relative encoding strength of color and form in a brain region. Because similarity within a feature changes across brain regions (e.g., two similar colors in one region may become dissimilar in another region), it would not have been possible to equate color and form variations for all the brain regions examined. Thus we have chosen what we believe to be reasonably large variations within each feature, including choosing two spirals with opposite directions, two mid-level form features with straight vs curved edges, and two hues that are maximally distinctive. These feature variations allow us to make a reasonable evaluation of the relative coding strength of color and form in each brain region, and more importantly, how the feature coding bias may change across visual regions.

Although it could be argued that perhaps a wider array of colors and form features could have been sampled, by using a small number of stimuli chosen to greatly differ with respect to a chosen dimension (hue, orientation, curvature), we were able to maximize our power, giving us more confidence that any null results were not due to an inadequate number of trials. Furthermore, the logic of the double-conjunction design we used requires two pairs of stimuli that differ with respect to two features.

To conclude, our comprehensive approach illuminates the overall architecture of color and form processing in the human brain. Color and form information was not anatomically segregated into distinct anatomical regions defined by their univariate sensitivity to either feature, but instead was generally co-localized in the same brain regions in a biased distributed manner throughout the ventral visual processing pathway, with decoding from color and shape regions largely consistent with their univariate preferences. This challenges a strictly modular view of color and form processing. Meanwhile, despite the co-existence of color and form information in most regions we examined, this joint coding was nearly always additive, with explicit encoding for specific *conjunctions* of these features occurring only for the conjunction of color with simple form features in early visual cortex. Thus, the predominant relationship between color and form processing in the human ventral visual hierarchy appears to be one of anatomical coexistence, but representational independence.

## Acknowledgements

This research was supported by a National Science Foundation Graduate Research Fellowship (DGE1745303) to J.T., a National Institute of Health Grant (1R01EY030854) to Y.X., and a NIH Shared Instrumentation Grant to Harvard Center for Brain Science (S10OD020039). We thank Talia Konkle, Geoge Alvarez, Alfonso Caramazza, and members of the Harvard Vision Lab and the Harvard Cognitive Neuropsychology Labfor their valuable feedback on this project.

---

**Supplemental Figure 1.**
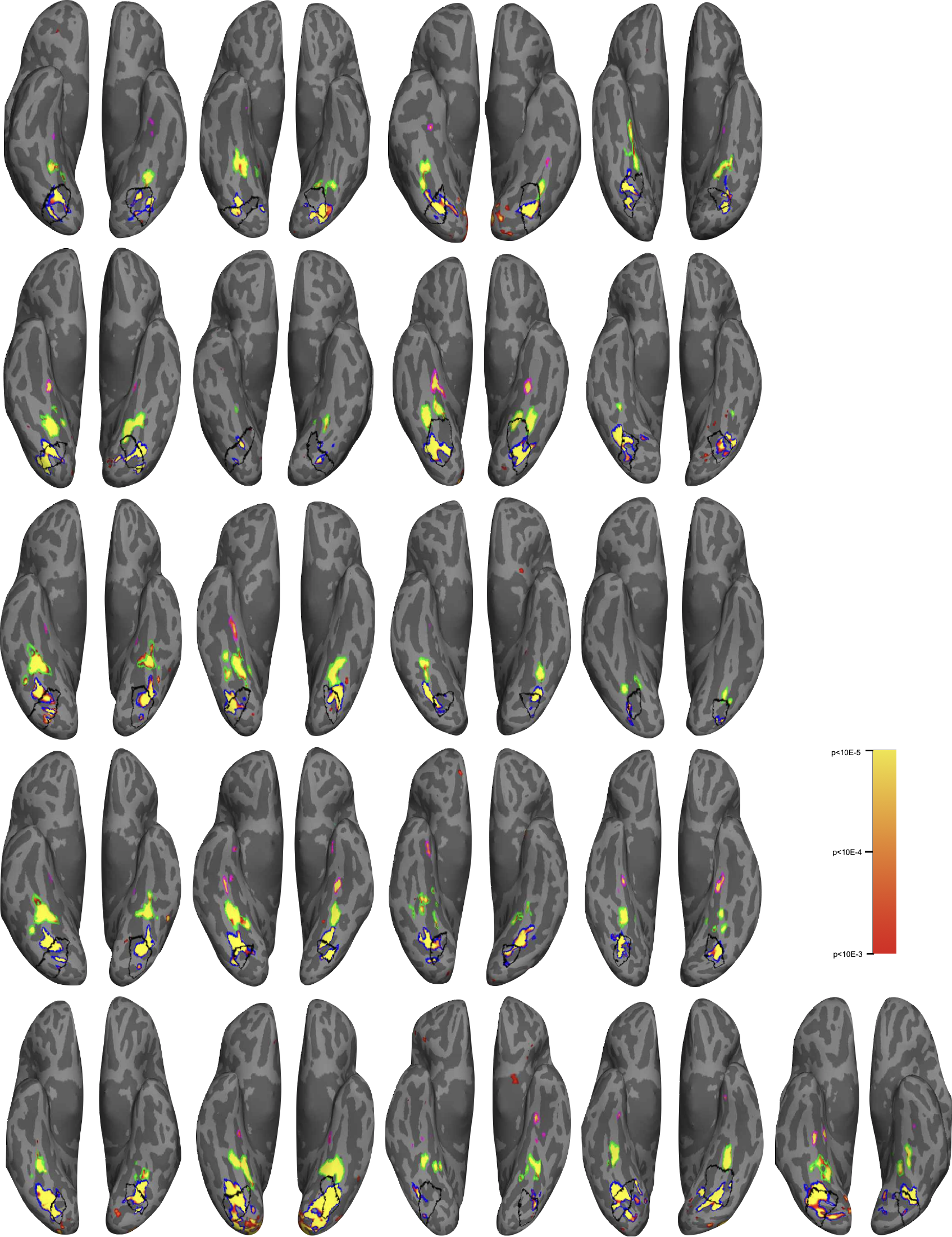
Ventral view of the brain showing the color regions for all participants across the two experiments defined as higher activation for color than greyscale scenes. The posterior, central, and anterior color regions and retinotopic V4 are shown in blue, green, magenta and black outlines, respectively.

